# Dynamic changes in the plasmidome and resistome in the gastrointestinal tract of chickens

**DOI:** 10.1101/2025.05.29.656840

**Authors:** Marketa Rysava, Jana Palkovicova, Katarina Stredanska, Jana Schwarzerova, Marketa Jakubickova, Darina Cejkova, Derya Aytan-Aktug, Saria Otani, Monika Dolejska

## Abstract

The expansion of intensive poultry farming has led to a substantial increase in antibiotic use, which in turn has promoted the accumulation of antibiotic resistance genes (ARGs). The chicken gut serves as a reservoir for these genes and provides favorable conditions for their horizontal transfer via mobile genetic elements, such as plasmids. Through this process, commensal bacteria can transfer ARGs to pathogens, facilitating their spread and increasing the risk of transmission to humans.

In this study, long-read sequencing was used to characterize plasmidome and resistome in 12 fecal samples from three houses of a commercial chicken broiler farm. All chickens received enrofloxacin in the first days of life, with one house additionally treated with sulfamethoxazole/trimethoprim combination. For comparison, metagenomic analysis using short-read sequencing was performed on the same samples.

This study revealed the presence of various ARGs associated with resistance to 26 antibiotic classes. Strong genetic association between MOBP-type plasmids and fluoroquinolone resistance was observed within chicken broiler farm. Temporal trends indicated progressive mobilization of these ARGs, suggesting an increasing potential for horizontal gene transfer. While fluoroquinolone resistance expanded over time, diaminopyrimidine resistance remained stable despite the antibiotic treatment. Most ARGs were carried on small plasmids, and complete plasmid reconstructions ranged from 2.6 to 47.6 kb.

Despite technical limitations, our findings demonstrate that plasmidome sequencing can enrich metagenomic analysis by enabling the detection of low-abundance plasmid types and providing deeper insights into the dynamic plasmid-mediated dissemination of ARGs in the chicken gut microbiome.

**Importance:** Despite the crucial role of plasmids in antimicrobial resistance (AMR) dissemination, studies focusing on plasmidome, defined as the complete set of plasmids, remain limited. Combining a metagenomic approach with a focus on plasmids enhances our ability to understand the genetic context and mechanisms underlying AMR transmission. The findings emphasize the importance of targeted plasmid analysis to improve surveillance and risk assessment of AMR transmission in microbial ecosystems.

## Introduction

Intensive poultry farming has led to increased antibiotic use, driving the rise of antimicrobial resistance (AMR), which is a major global health concern. Despite the prohibition of antibiotics as growth promoters in EU livestock farming since 2006 and a ban on their use for disease prevention in 2022, more than half of global antibiotic consumption is still attributed to agriculture (1). In the Czech Republic, penicillins, tetracyclines and sulfonamides are the most commonly administered to livestock (2). While fluoroquinolones are not among the top antibiotic classes by total weight, they are the most frequently administered antimicrobial group in the poultry sector which account for almost 80% of fluoroquinolone consumption in livestock. Although sales of fluoroquinolones fluctuated between 2011 and 2022, there was a slight overall increase from 1.5 mg/Population Correction Unit (PCU) in 2011 to 1.6 mg/PCU in 2022. Nevertheless, during the implementation of the second Czech national action plan on antimicrobial resistance (2019–2022), fluoroquinolone consumption decreased by 13%. (2, 3). In contrast, fluoroquinolones are not approved in poultry farming in the United States (4) and Australia (5).

The gut microbial community benefits the host through competitive exclusion, preventing pathogen colonization by outcompeting them for nutrients and adhesion sites. It also plays a crucial role in modulating immune responses, enhancing pathogen clearance, and supporting host nutrition through metabolic byproducts that promote gut health. However, despite the beneficial roles, the gastrointestinal microbiome can also serve as a reservoir of antibiotic resistance genes (ARGs). These genes may be transferred between bacteria via horizontal gene transfer (HGT), further facilitating the persistence and spread of AMR within the host and its surroundings (6). The resistant bacteria originating from the poultry gut may be transmitted to humans through the food chain, particularly via contaminated chicken meat (7) or introduced into the environment through the application of chicken manure as fertilizer (8). Approximately 60% of human infectious diseases are estimated to be caused by zoonotic pathogens capable of carrying ARGs (8).

Plasmids play a central role in HGT (9). They are characterized by significant variability, their size ranging from a few kilobases to hundreds of kilobases, and their copy number differing markedly between cells. Despite this characteristic, plasmid DNA (pDNA) constitutes only a small fraction of the total cellular DNA compared to chromosomal DNA (10, 11). The plasmidome, defined as the total collection of plasmids in a specific environment, plays a significant role in microbial evolution and adaptation (12).

The number of studies focusing on the plasmidome, particularly in the context of the chicken gut, is limited. Previous publications were focused on the plasmidome in sewage (12–15) or in the gut of other animals (16, 17), however, the role of plasmidome in the spread of AMR in chicken gut remains largely unexplored. The focus of this research was to characterize plasmidome and resistome in the fecal droppings of chickens, as well as to obtain complete plasmid sequences and link them to specific resistance genes using long-read sequencing technology. This was followed by tracing the temporal changes in the plasmidome to investigate the dynamics and evolution of plasmid-mediated AMR during the broiler chicken’s lifetime in respect to the use of antibiotics.

## Materials and Methods

### Study design and sampling

Twelve samples of fecal droppings were obtained from broiler Ross 308 flocks located in three houses. From each house, sampling began in the second week of life and continued weekly for three weeks until the birds reached slaughter age. Only fecal material was collected at designated time points, with no direct involvement of the animals (detailed ethical statement in Supplementary material S1). Chickens in all houses were treated with enrofloxacin in the first days of life. One of the houses was subsequently treated with a sulfamethoxazole/trimethoprim combination due to higher mortality. A composite fecal sample from each house was taken each week. The uric acid layer was manually removed from the chicken droppings to prevent potential inhibition of DNA extraction, and each sample was stored at −20°C.

### DNA extraction and post-extraction treatment for long-read sequencing

Total pDNA was extracted from the obtained samples using Plasmid Mini Kit (Qiagen, GE) and Plasmid Midi Kit (Qiagen, GE) to obtain both small and large plasmids. The extractions were performed in accordance with the manufacturer’s guidelines with modifications provided in Supplementary material S1. After extraction, residual chromosomal and environmental DNA were removed using Plasmid-Safe ATP-Dependent DNase (10 U) (Biosearch Technologies, UK), and pDNA was subsequently amplified using NxGen phi29 DNA Polymerase (10 U) (Biosearch Technologies, UK). Detailed protocol for post-extraction treatment is provided in Supplementary material S1. Treated pDNA from the Plasmid Mini Kit and Plasmid Midi Kit was mixed 1:1 for each sample.

### DNA extraction for short-read sequencing

A total of 250 mg of droppings was used for DNA purification using the QIAamp® PowerFecal® Pro DNA Kit (QIAGEN, GE), following the manufacturer’s protocol with minor modifications. Sample homogenization was carried out using the bead-beating step provided in the kit, employing the Vortex-Genie 2 and horizontal adapter (Scientific Industries, USA). DNA was eluted following a 5 min incubation with the provided elution buffer. DNA yield was quantified using a NanoPhotometer N60 (Implen, GE), and DNA integrity was assessed by running a 1% agarose gel.

### Plasmidome and Metagenome Sequencing

For long-read sequencing of plasmidome, DNA libraries of each sample were prepared from the treated pDNA using the SQK-LSK114 ligation kit (Oxford Nanopore Technologies, ONT, UK). The manufacturer’s guidelines were followed, with modifications specified in Supplementary material S1. Each DNA library was loaded onto a FLO-PRO114M flow cell, one sample per flow cell, and sequenced on the P2Solo long-read sequencing platform (ONT, UK).

For short-read sequencing, DNA was diluted to a concentration of 10 ng/µL. DNA libraries were prepared from 10 ng of metagenomic DNA using VAHTS Universal Plus DNA Library Prep Kit for Illumina (Vazyme Biotech, China) and sequenced via paired-end chemistry (PE150) on the NovaSeq X platform (Illumina, USA) by Biomarker Technologies Co., Ltd. (Beijing, China). The average yield per sample was 12.03 Gb of sequencing data, with the percentage of Q30 bases in each sample exceeding 91.89%.

### Long-read data analysis

Raw data obtained from long-read sequencing was pre-processed and GraphPad Prism v10.0.0 (GraphPad Software, Boston, USA) was used for subsequent data comparison (Supplementary material S1). The statistical analysis was performed using Pearson correlation, with a significance threshold set at *p* ≤ 0.05. To compare sample libraries treated with short fragment buffer (SFB) and long fragment buffer (LFB), the FastQC statistics, assembly statistics and the results obtained by PPR-meta tool v1.1 (18) were visualized in Python using Matplotlib library v3.4.4 (19).

KMA tool v1.4.15 (20) with option ‘bcNano’ was used to compare the genetic content of processed reads. The ARGs were assessed using PanRes v1.0.1 database (21). The mobile genetic elements (MGEs), besides plasmids, were identified by the MGE v1.0.2 database (22) and the plasmid content was analyzed by mapping to the *oriT* database of the MOB-suite v3.1.9 (23). To obtain comparable results across different samples, the relative abundance was calculated and normalized using Reads Per Kilobase Million (RPKM) value. Twenty most abundant MGEs and plasmids were categorized as distinct groups while the remaining elements were consolidated into an ‘others’ group. The resulting data were visualized using the Matplotlib library v3.4.4 in Python.

To further investigate the co-occurrence of MGEs and ARGs across houses and time points, a network-based approach was employed (24). Seven datasets (Table 1) were generated from the alignment files produced by the KMA tool for ARGs and plasmids.

**Table 1.**
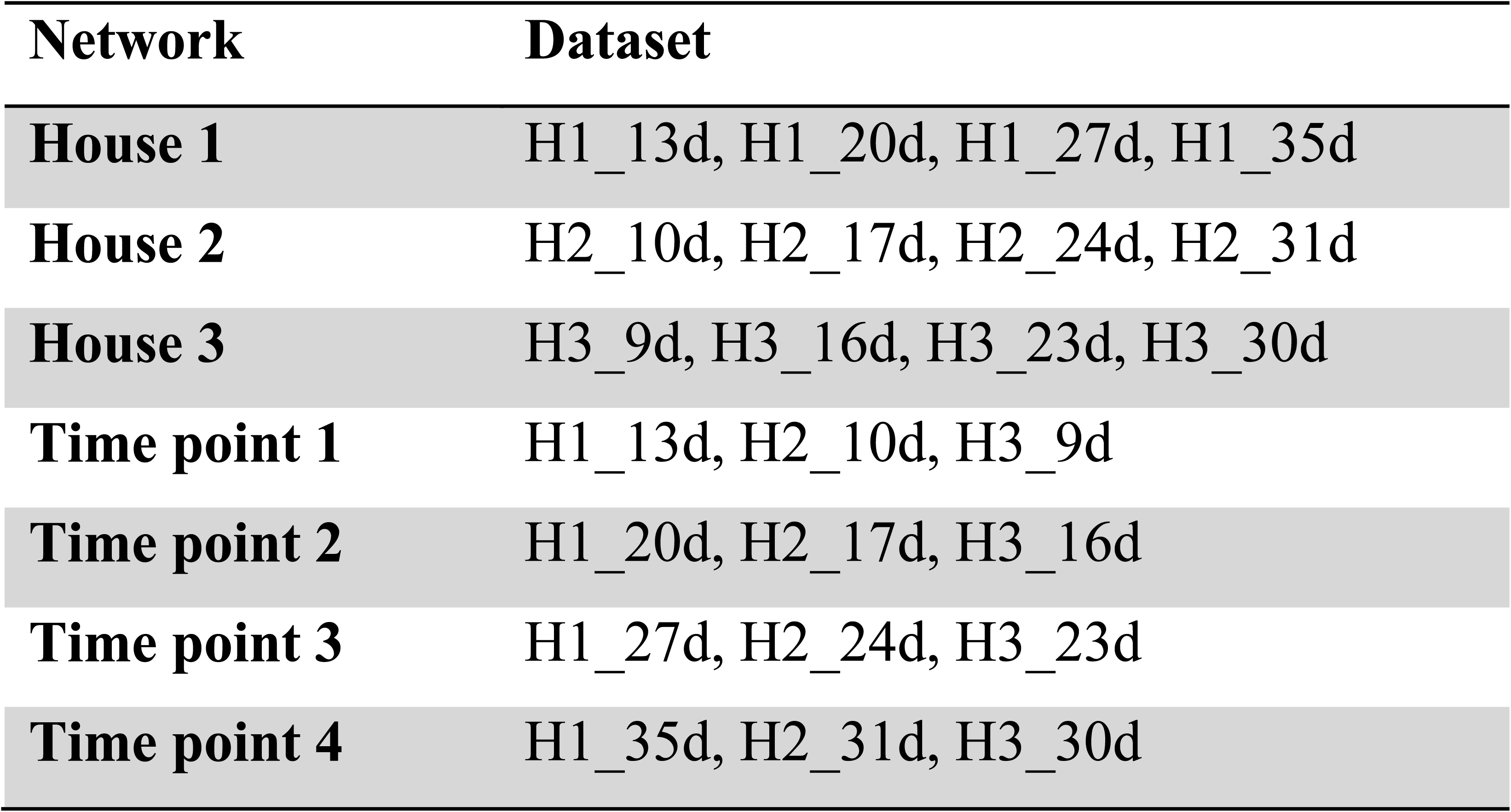
Datasets used for network analysis.

Count-transfer-matrix, obtained as described (Supplementary material S1), was used for the construction of an undirected network which was visualized in Cytoscape v3.9.0 (25). As the network represents node co-occurrence and not their similarity, node positions were manually adjusted for improved clarity. However, the network retains the original connectivity.

Plasmids carrying ARGs were assembled and corrected (Supplementary material S1). The assemblies were analyzed using the reconstruction mode of MOB-suite (23). The ARGs were identified by comparing the assemblies to the PanRes v1.0.1 database using ABRicate (https://github.com/tseemann/abricate) with the same criteria as in the raw read analysis. Plasmids carrying ARGs were compared with PLSDB v. 2024_05_31_v2 (26) [accessed on November 2024] and annotated using Bakta v1.10.3 (27) with subsequent manual curation in Geneious Prime® 2024.0.7 (http://www.geneious.com/).

### Short-read data analysis

Short metagenomic sequencing reads were pre-processed (Supplementary material S1). Trimmed, contamination-free sequencing reads were used for taxonomic profiling with MetaPhlAn v4.0.6 (28) and the embedded GTDB release 207 (29). Genetic content (ARGs, MGEs and plasmids) of the processed reads was assessed using the KMA tool v1.4.15 with option ‘1t1’ and databases PanRes v1.0.1, MGE v1.0.2 and MOB-suite v3.1.9 with subsequent analysis as described for long-read processing of genetic content.

## Results and discussion

### Plasmid DNA extraction and library preparation efficiency

Total pDNA extracted from the twelve chicken waste samples from three different houses of a chicken farm in the Czech Republic yielded concentrations ranging from 1.03 ng/µL to 3.4 ng/µL using Plasmid Mini Kit and 1.12 ng/µL to 2.74 ng/µL using Plasmid Midi Kit (Table S1). DNA extraction from chicken droppings is particularly challenging due to the presence of various inhibitors and the relatively low concentration of bacterial DNA in the samples (30). An even greater challenge is the extraction of plasmids, particularly for low-copy-number plasmids or those with minimal plasmid content relative to chromosomal DNA (12). Following DNase treatment to remove chromosomal DNA, the reduction in pDNA concentration varied between 60.00% and 95.15% for the pDNA extracted by Plasmid Mini Kit, and between 37.12% and 87.50% for the Plasmid Midi Kit. Subsequent amplification process yielded pDNA concentrations ranging from 14.9 ng/µL to 94.3 ng/µL for the Plasmid Mini Kit and between 18.7 ng/µL and 106.0 ng/µL for the Plasmid Midi Kit, providing sufficient material for sequencing. Library preparation resulted in reduced concentration following LFB washing, with reductions ranging from 22.77% to 94.95% across samples. However, when compared to SFB, which does not remove small fragments (<3 kb), the reduction remained comparable for the tested sample (H1_27d). The H1_27d had an initial concentration of 38 ng/µL pre-wash, decreasing to 28.4 ng/µL after washing with SFB and to 27 ng/µL with LFB. Washing with SFB buffer resulted in an increased number of fragments below 1000 bp and, more importantly, a higher proportion of residual chromosomal DNA (Fig. S1). In this study, we optimized DNA purification strategies (use of two plasmid DNA isolation kits, pre-treatment with SDS and lysozyme, buffer volume adjustment, extended times, amplification in triplicates). Additionally, linear DNA digestion and circular element enrichment steps were employed to increase the plasmidome yield. These optimizations improved plasmid yields, particularly for plasmids from diverse bacterial species or complex microbial communities. However, further refinement would be needed for routine experiments.

### Association of pore count and data quality

Statistical analyses revealed the relationships between DNA concentration, the number of flow cell pores, sequencing output, read lengths, and the lengths of assembled contigs (Fig. 1). Pearson correlation analysis revealed a weak positive relationship between the number of pores on the flow cell and the number of generated bases (*r* = 0.337, *p* = 0.286). While this suggests a tendency for a higher pore count to result in increased base output, the correlation was not statistically significant.

**Fig. 1:**
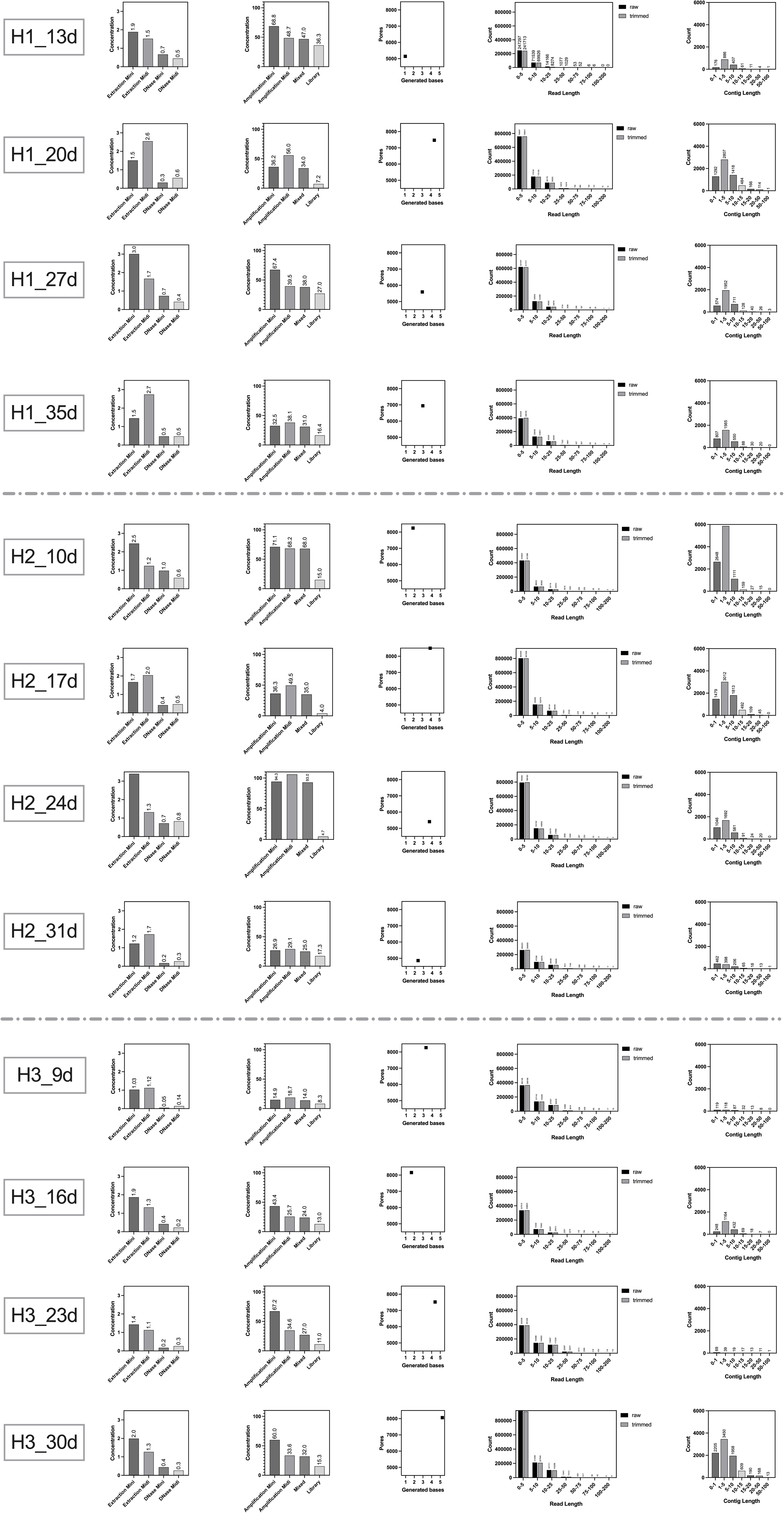
Overview of quality control and sequencing statistics for plasmidome samples across the chicken farm. Each row represents one sample collected from a specific house and time point. The columns show (from left to right): DNA concentration after extraction and DNAse treatment (ng/μL), DNA concentration after amplification and library preparation (ng/μL), number of generated bases (Gb), read length (kb) distribution (raw vs. trimmed), and contig length (kb) distribution after assembly.

A highly significant, strong linear relationship exists between the amount of generated data and total read length, both with raw untrimmed (r = 0.9979, *p* = 0.00001) and trimmed (r = 0.9983, *p* = 0.00001) sequencing data. This confirms that a higher number of generated bases corresponds to longer reads, suggesting that increasing sequencing output leads to an overall increase in read length. In contrast, the number of generated bases did not significantly affect either the number of contigs (*r* = 0.28, *p* = 0.38) or their total length (*r* = 0.4573, *p* = 0.13). Similarly, total read length had no significant impact on contig length (*r* = 0.48, *p* = 0.11) or the number of contigs (*r* = 0.291, *p* = 0.36). All data is available in the supplementary material (Table S1).

### Mobilome

#### Mobile genetic elements identified via long-read sequencing

The analysis of MGEs, excluding plasmids, showed a wide variety of identified elements, including miniature inverted-repeat transposable element (MITE), insertion sequence (IS), mobilizable transposon (MTn) and transposon (Tn) (Fig. S2, Table S2).

In the plasmidome, IS*Ljo1* and IS*Lre2* appeared at all time points across all three houses. Similarly, MITE*Ec1* was present in the majority of the time points. House 1 was characterized by the high prevalence of the MITE*Ec1* and IS*Ljo1* over time, with different elements dominating at various time points. In house 2, there was greater variability in MGE representation, with elements such as IS*Lre2* and IS*Lhe30* being notable. House 3 exhibits the most pronounced differences in MGE representation, with IS*Enfa4* and IS*679* showing high RPKM values during the first and last time points (Fig. S2 A).

The proportion of MGE groups in the metagenome is comparable to that observed in the plasmidome. The most abundant occurring insertion sequence is IS*Ljo1*, which, similarly to the plasmidome, is present at all time points and in all houses. However, MITE*Ec1*, IS*Enfa4* and IS*679* exhibit lower abundance in metagenome. In general, there were no significant differences in the total RPKM values between the metagenome and plasmidome, indicating that MGEs detected in total metagenomic samples are not confined solely to plasmids but may also reside on chromosomal DNA (Fig. S2 B).

#### Plasmids

Plasmids detected in the plasmidome analysis exhibited significantly higher variability compared to metagenomic analysis (Fig. 2 A, C). Across the entire plasmidome dataset, a total of 133 diverse plasmid types were detected, with the number of plasmid types ranging from 20 to 75 per sample. In comparison, the metagenome comprised 59 unique plasmid types, with per-sample diversity ranging from 1 to 31 (Table S2). Notably, single plasmid (NZ_CP009579|MOBV) was detected in the H3_30d metagenome (Fig. 2D), whereas the corresponding plasmidome contained a total of 50 unique plasmid types (Table S2). Temporal shifts in plasmid composition revealed that certain plasmids, such as KY014464|MOBP and JN985534|MOB_unknown, persisted across samples in plasmidome and were also detected in the metagenome (Fig. 2 B, D).

**Fig. 2:**
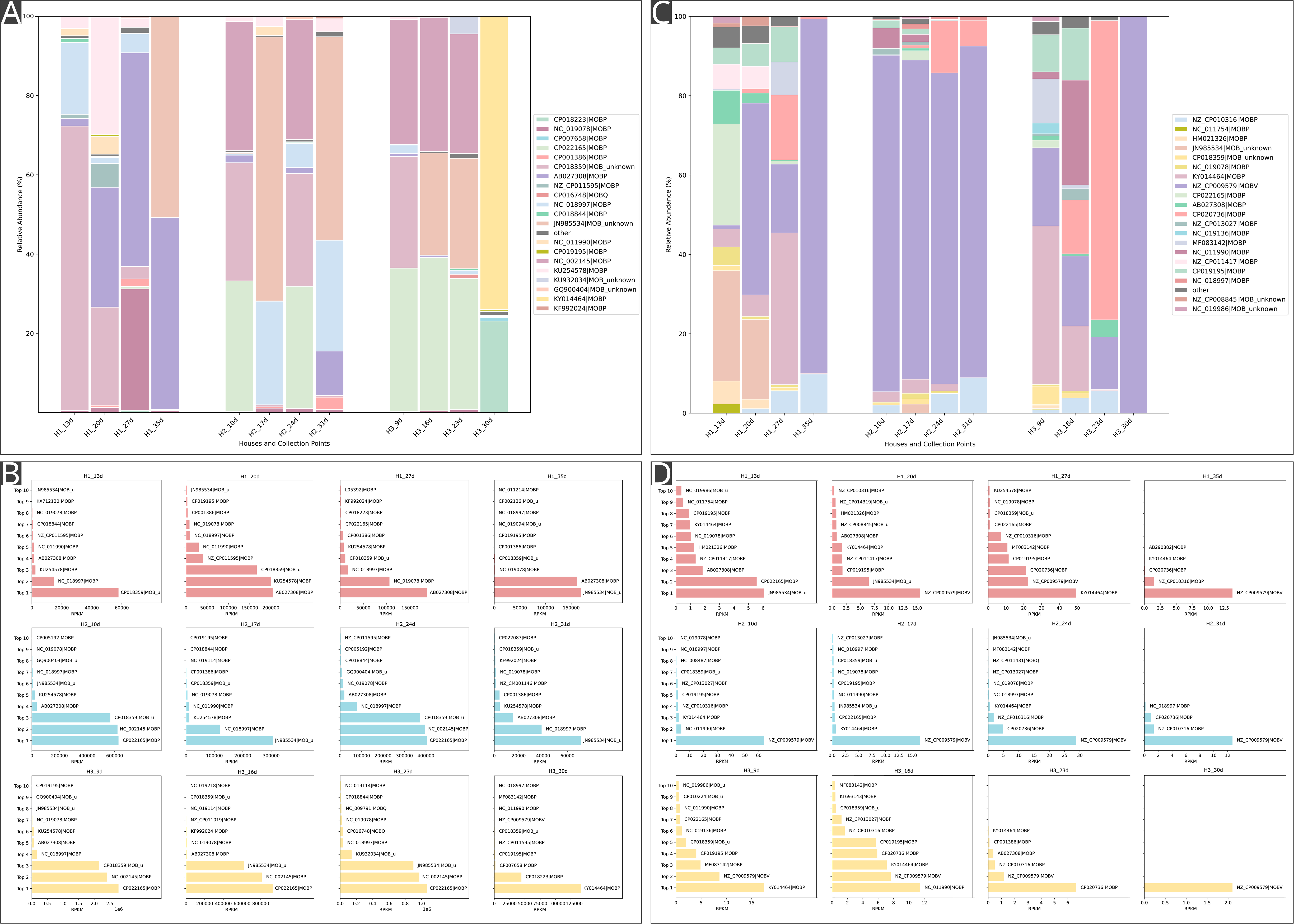
Relative abundance and distribution of plasmid types across different sampling points and houses. The study was performed on a chicken farm comprising three separate houses (H1, H2, H3), each sampled at four different time points. Stacked bar plots show the relative abundance of various plasmid types in the plasmidome (A) and metagenome (C) fractions of the three chicken houses (H1, H2, H3), each sampled at four different time points. Because of the plasmid diversity, each bar plot has its own color code specified in the legend. The top 20 plasmid groups were categorized separately, remaining plasmids were grouped as ‘others’. Horizontal bar charts present the top 10 most abundant plasmid types identified in each sample from the plasmidome (B) and metagenome (D).

In general, plasmids show significantly lower abundance in metagenomic datasets compared to their representation in the plasmidome (Fig. S3, Table S1). The highest total RPKM value among all samples in the metagenome is 129.48, whereas the lowest value in the plasmidome is 80,503.15. The amplification step using phi29 polymerase during plasmidome preparation likely affected plasmid abundance, potentially skewing quantitative comparisons. In contrast, the treatment with Plasmid-Safe DNase, intended to remove linear DNA, may have influenced plasmid diversity by inadvertently degrading fragmented large plasmids. Nevertheless, long-read plasmidome sequencing enabled the detection of low-abundance plasmid types that would otherwise remain undetected. These observations underscore the importance of not only filtering out chromosomal sequences prior to plasmid analysis but also extracting plasmids separately as essential steps for their comprehensive characterization. Although the metagenomic approach (8, 31) provides valuable insights into complex microbiomes including unculturable segments of bacterial communities, plasmids and other MGEs are not adequately sequenced due to their chromosome size (32). Plasmidome analysis (12, 16, 17) allows for more accurate detection of plasmids and thus better assessment of HGT.

### Antibiotic resistance genes

The identified ARGs in plasmidome and metagenome were associated with 26 antibiotic groups (Fig. 3A, Table S2).

**Fig. 3:**
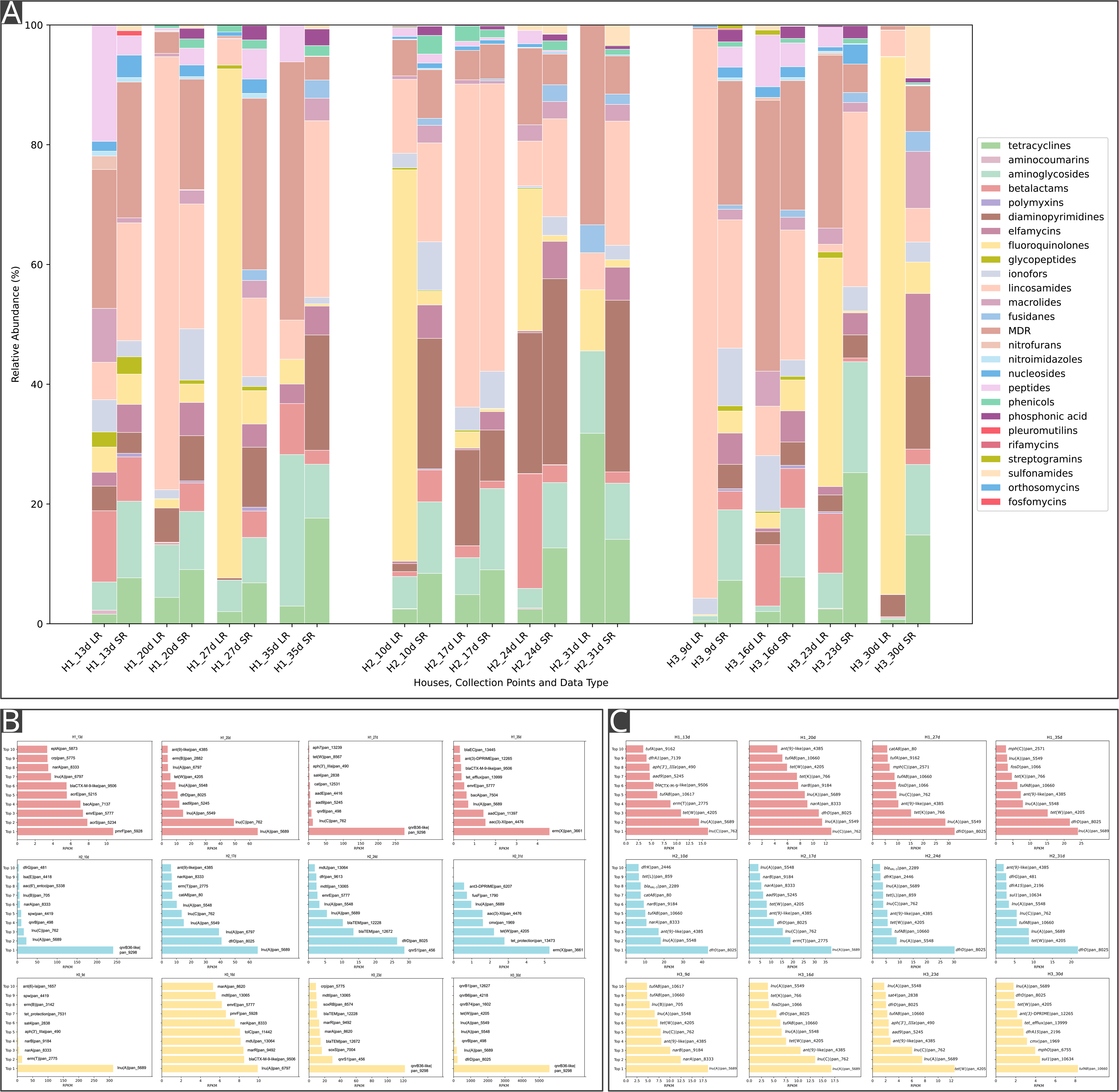
Relative abundance and distribution of ARGs across different sampling points and houses. Stacked bar plot (A) displays differences in relative abundance of genes conferring resistance to 26 AMR groups in the three houses (H1, H2, H3) of a commercial chicken farm in Czech Republic collected within four time points between two datasets: the plasmidome, obtained by long-read sequencing (LR), and the metagenome, obtained by short-read sequencing (SR). Horizontal bar charts present the top 10 most abundant AMR genes identified in each sample from the plasmidome (B) and metagenome (C). MDR - multidrug resistance

A comparison of the plasmidome and the metagenome revealed notable differences in the representation of ARGs. The relative abundance of fluoroquinolone resistance genes was significantly lower in the metagenome, suggesting that these genes are primarily carried by plasmids. Certain resistance groups, such as aminocoumarins, were absent in the metagenome, indicating that these genes are likely plasmid-encoded as well. Conversely, the orthosomycins and fosfomycin groups did not appear in the plasmidome, which implies that these genes are more likely located on the chromosome in our dataset. (Fig. 3A).

The variants of *qnr* for fluoroquinolone resistance were the most frequently found genes in plasmidome. The *qnrB36*-like|pan_9298, which was highly abundant in sample H3_30d (5597.9 RPKM), was identified among the plasmidome, with high abundance in samples H1_27d, H2_10d, and H3_23d. In contrast, this gene was not among the top 10 most abundant genes in the metagenome (Fig. 3 B, C). All houses were treated with enrofloxacin during the early days of life, which may be a reason for an expansion of genes conferring resistance to fluoroquinolones.

House 2 was subsequently treated with a combination of sulfamethoxazole and trimethoprim. This therapeutic intervention could explain the observed increase of diaminopyrimidine resistance genes in the plasmidome between the first and third time points in house 2. This trend did not persist until the time point four, where diaminopyrimidine resistance genes were no longer detectable. However, in the metagenome, diaminopyrimidine resistance genes were stable in all houses. These findings may indicate that resistance to fluoroquinolones and diaminopyrimidines could be plasmid-associated. Notably, sulfonamide resistance genes remained at low or undetectable levels both in the plasmidome and the metagenome. In other studies, tetracyclines frequently rank among the most detected ARGs in chicken manure, which is consistent with their widespread use in livestock farming, where they are commonly administered in significant quantities (8, 31, 33). Contrary to the previous studies, tetracycline resistance genes were not as prominently represented in our samples compared to other resistance groups. For example, lincosamide resistance genes exhibited high abundance in both metagenome and plasmidome. In our sample set, variants of *lnu*(A) were a top 1 gene with the highest abundance among different metagenomic samples (H1_13d, H1_20d, H1_35d, H2_17d, H3_9d, and H3_16d) as well as plasmidomic samples (H1_20d, H2_17d, H3_9d, and H3_16d) (Fig. 3 B, C). This suggests their presence on both plasmids and chromosomes. In a previous study, *lnu*(A) was the gene with the highest abundance in chicken litter (8).

Our findings are consistent with previous studies and suggest that antibiotic administration contributes to the selection and prevalence of resistance genes (34, 35).

### Plasmids associated with antibiotic resistance genes

Using network-based co-occurrence analysis, genetic associations between particular plasmids and ARGs were detected to reveal their potential role in ARG dissemination (Fig. S4).

Across all three houses, the strongest association was consistently observed between MOBP group plasmids and fluoroquinolone resistance genes (Fig. 4). Among these, plasmid CP018223|MOBP showed the most prominent and consistent co-localization with *qnrB46*-like|pan_9298 (20,712 reads), suggesting that the gene is likely carried by this plasmid. A notable co-occurrence was also detected with *qnrB*|pan_498. In house three, CP018223|MOBP further co-occurred with *qnrB6*|pan_4218 and *qnrB74*|pan_1602, although these associations appeared comparatively weaker (Table S3).

**Fig. 4:**
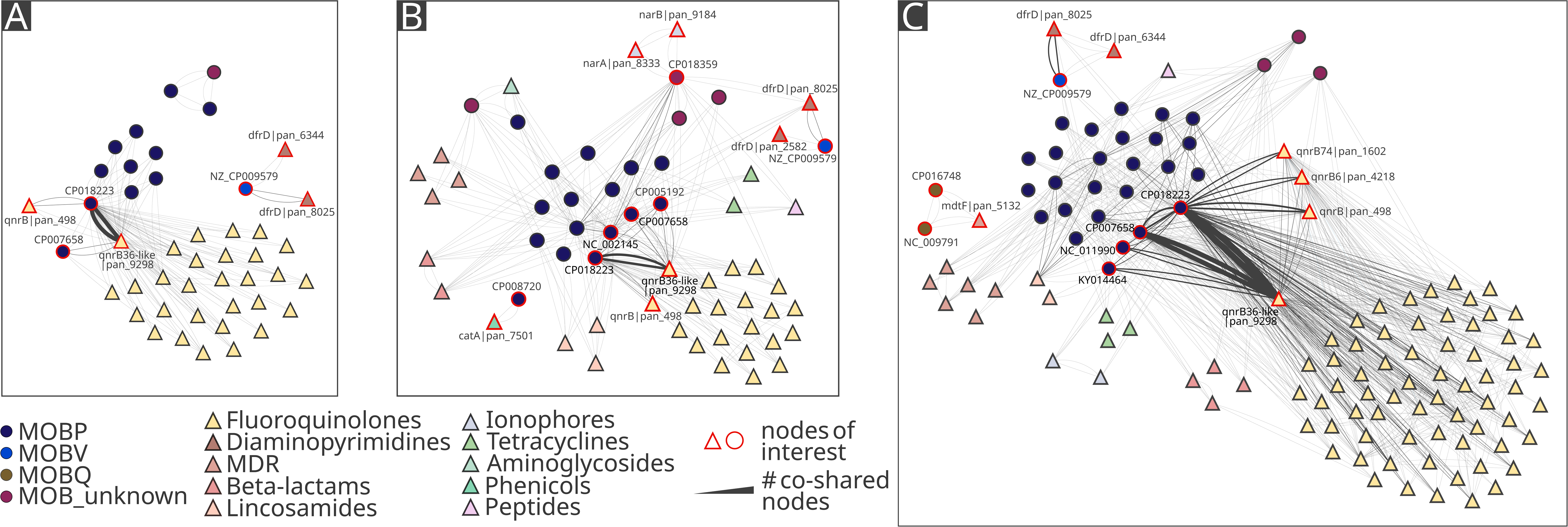
Co-occurrence network of antibiotic resistance genes and plasmids in different houses. The undirected network, where nodes represent plasmids (circles) or antibiotic resistance genes (triangles), and edges represent their co-occurrence on the same sequence. The width of the edges corresponds to the number of sequences co-sharing the specific nodes ranging from one read to 20,712 reads. Layout does not represent similarity or spatial distance as node positions were manually adjusted for an improved visual clarity.

Furthermore, the most frequently occurring gene, *qnrB46*-like|pan_9298, was also likely present on plasmid CP007658|MOBP across all houses, although at a much lower frequency. In house two, *qnrB46*-like|pan_9298 was associated with plasmids NC_002145|MOBP and CP005192|MOBP, while in house three, it was found to co-occur with NC_011990|MOBP and KY014464|MOBP, although less frequently.

The NZ_CP009579|MOBV plasmid was most frequently associated with genes encoding resistance to diaminopyrimidine antibiotics across all houses. In house two, two genes encoding resistance to ionophores co-localized with plasmid CP018359|MOB_unknown. Additionally, this house also contained the *catA*|pan_7501 associated with plasmid CP008720|MOBP. In house three, two MOBQ plasmids (NC_009791|MOBQ and CP016748|MOBQ) were associated with the *mdtF*|pan_5132 gene, which provides MDR. Overall, the associations found between resistance genes and plasmids suggest that plasmids may play a key role in the spread of resistance genes to the following classes of antibiotics: fluoroquinolones, diaminopyrimidines, MDR, beta-lactams, lincosamides, ionophores, tetracyclines, aminoglycosides, phenicols, and peptides (Fig. 4).

The longitudinal analysis of plasmid-ARGs associations indicates that fluoroquinolone resistance genes increased notably in both frequency and diversity of plasmid linkages. At the early time points, these genes were present but showed only moderate plasmid connections (Table S3). However, their interactions expanded substantially in the following weeks, involving a broader range of genes and plasmids (Fig. 5). Our observations correspond to the previously reported findings (36), which demonstrate that even a single course of enrofloxacin treatment can directly contribute to both, the emergence and persistence of fluoroquinolone resistance in *Campylobacter coli*.

**Fig. 5:**
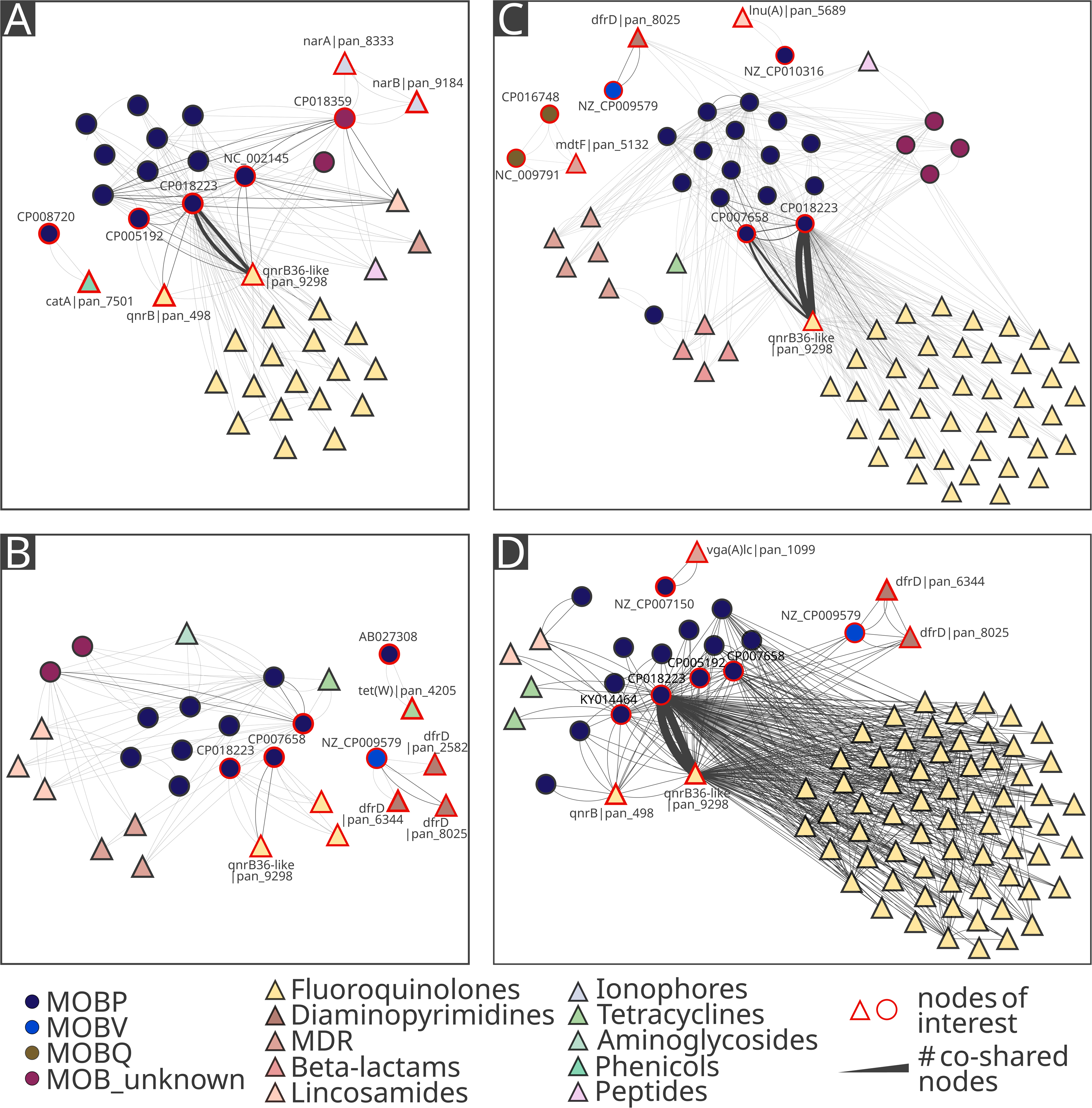
Co-occurrence network of antibiotic resistance genes and plasmids in different time points. The undirected network, where nodes represent plasmids (circles) or antibiotic resistance genes (triangles), and edges represent their co-occurrence on the same sequence. The width of the edges corresponds to the number of sequences co-sharing the specific nodes ranging from one read to 20,253 reads. Layout does not represent similarity or spatial distance as node positions were manually adjusted for an improved visual clarity.

Genes conferring resistance to diaminopyrimidines exhibited relatively stable plasmid associations over time and across houses (Fig. 5). Despite the use of trimethoprim-containing treatment in the second chicken house, no substantial increase in *dfr* gene prevalence or diversity of plasmid associations was observed. This suggests that the treatment-induced selective pressure did not result in detectable mobilization or expansion of diaminopyrimidine resistance genes within the sampling. Our observations are not in concordance with previous findings reporting a resistance acquisition in *Staphylococcus aureus* strains exposed to sulfamethoxazole/trimethoprim treatment *in vitro* (37) or an increase in *Escherichia coli* resistance following this treatment in pigs (38).

The assembly of plasmidome samples produced a high number of contigs, ranging from 169 to 9818 per sample. We observed a high proportion of non-circular plasmid fragments as well as pieces of chromosomes. Frequently, the plasmid fragments appeared to represent duplicated plasmids within a single contig. This was supported not only by several repetitions of a single plasmid marker (e.g. plasmid replicon) but also by post-assembly mapping of corresponding reads. Based on the assembly analysis, it was revealed that some of the identified ARGs were localized on chromosomal fragments (Table S4).

Contrary to the previous research on the role of large plasmids on the dissemination of ARGs (39), the identified ARGs representing the chicken gut microbiome were mostly carried by small plasmids. This could be a result of the increased degradation potential of larger plasmids during the pDNA extraction, higher plasmid copy number of small plasmids and a higher potential of phi29 amplification of small plasmids. However, this could also be a result of microbiome composition which could lead to a plasmid loss due to secreted products from lactic acid bacterial genera from a class of Firmicutes present in the chicken gut microbiome (40).

Across all assembled samples, ARGs to 18 different groups of antibiotics were detected (Table S4). The MDR-conferring ARGs were strictly associated with chromosomal sequences while ARGs to lincosamides were present in both chromosomal and plasmid contigs. On the other hand, ARGs variants for fluoroquinolones were mostly linked to plasmids. We reconstructed five complete plasmids carrying ARGs ranging from 2.6 to 47.6 kb in size (Fig. 6). Many ARGs were localized on linear plasmid fragments which could not be fully reconstructed. Many candidate plasmids contained no ARGs. However, these can serve as a potential source for further dissemination of detected ARGs.

**Fig. 6:**
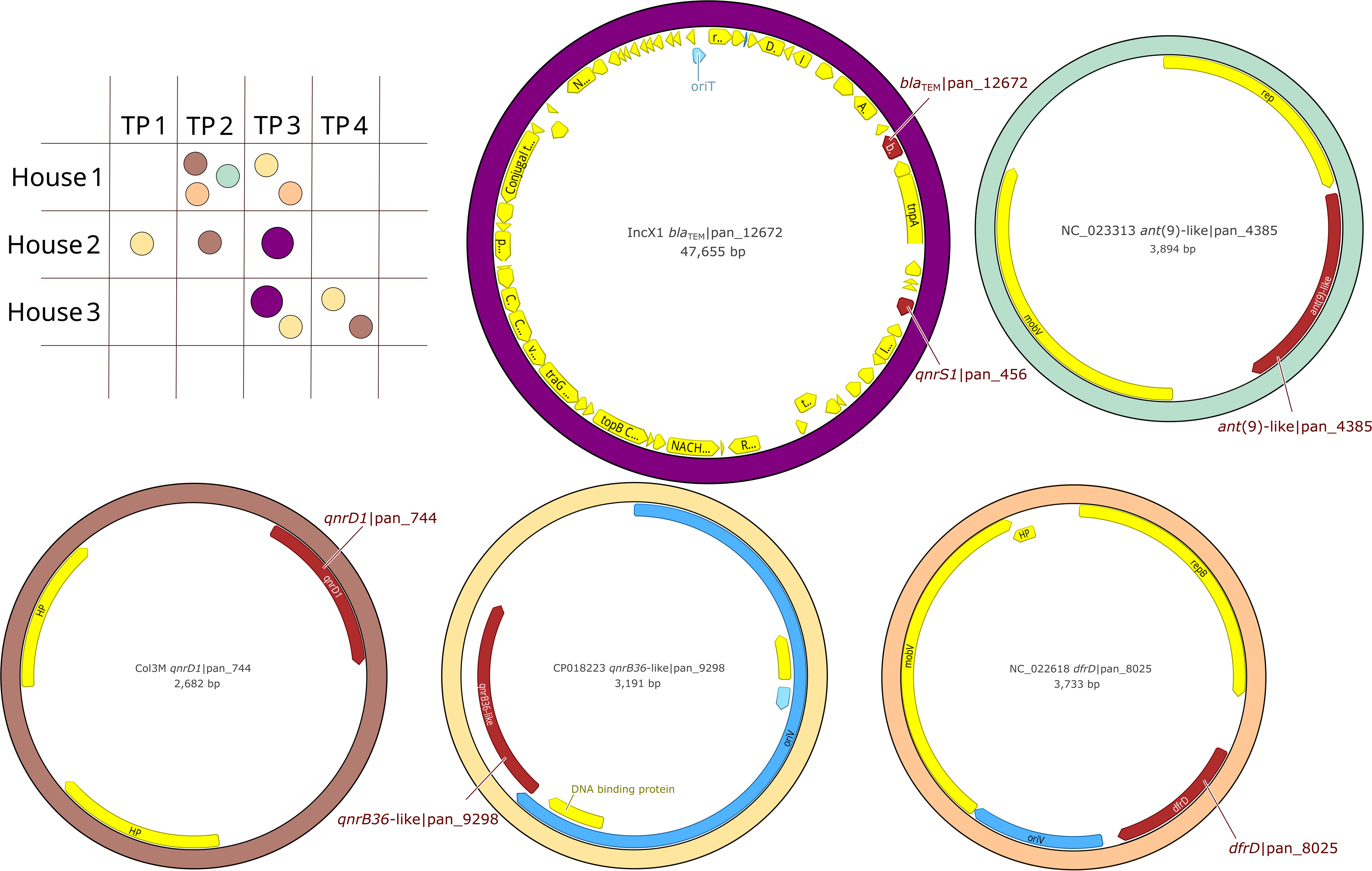
Complete plasmids associated with antibiotic resistance in samples from a chicken farm. The diagram (top left) indicates the presence of selected plasmids in individual samples collected from the three different houses (House 1–3) across four time points (TP1– TP4). Circular maps illustrate the annotated genetic structure of five representative plasmids assembled from long-read sequencing data, with antibiotic resistance genes (highlighted in dark red), origin of transfer (blue), and additional coding sequences (yellow).

In two (H1_35d, H2_31d) out of twelve samples, no ARGs were identified in assembled plasmids while in three other samples (H1_13d, H3_9d and H3_16d), all present ARGs were on chromosomal fragments (Table S4). The most occurring plasmid across the samples was a 3 kb long MOBP-like plasmid of rep_cluster_2335 containing *qnrB46*-like|pan_9298. This plasmid was found at different time points among all three houses (H1_27d, H2_10d, H3_23d and H3_30d), from the same predicted host (*E. coli*) with slight variations in size. The occurrence of this plasmid spiked in the H3_30d with coverage 88,543 which was also responsible for a huge increase in fluoroquinolone resistance observed in raw read plasmidome analysis. The 3,191 bp long plasmid from H3_30d is highly similar (99.96% similarity) with a previously published plasmid (OQ787038.1) from *E. coli* isolated from Eurasian coot and the 3,071 bp plasmid from H3_23d with plasmid (CP043749.1, similarity 99.96%) from *Salmonella enterica* isolated from pork.

Another ARG for fluoroquinolone resistance was *qnrD1*|pan_744 carried by a small 2.6 kb long Col3M plasmid present in H1_20d and H1_27d. The predicted host for these plasmids is *Proteus mirabilis*. The previously described plasmids (AP024494.1, CP057825.1) of highest similarity (99.96% and 99.90%, respectively) were associated with *Providencia vermicola* from human sample in Nepal and *Citrobacter freundii* from pig feces in United Kingdom. Therefore, these can be disseminated among diverse bacteria and circulated among humans and animals.

The *qnrS1*|pan_456 was co-harbored with *bla*_TEM_|pan_12672 on a 47.6 kb long IncX1 plasmid. This plasmid was found at the third collection point in house 2 and house 3 (H2_24d, H3_23d). These plasmids are highly similar (99.91%) to a previously described plasmid (MH121702.1) from Czech Republic found in rook feces. However, these plasmids were also identified in human bloodstream infections, pig feces and broiler samples (CP115825.1, CP122865.1, LR882060.1) from diverse countries isolated in different years. All of these plasmids are associated with *E. coli* strains.

The plasmids containing *dfrD*|pan_8025 were identified in H1_20d and H3_30d. However, from the raw read analysis, it is apparent that the gene was also present in house 2 (H2_17d). The most similar plasmid was from *Bacillus subtilis* which is 8.7 kb long and contains *tet*(M) and *tet*(L) genes instead of *dfrD*. The plasmids are 100% identical with coverage 80% which corresponds to the difference in the ARGs content.

The last fully assembled plasmid was 3.9 kb long, with replicon rep_cluster_1118 of MOBV group originating from *Staphylococcus aureus* which carried *ant*(*9*)-like|pan_4385, also known as a gene *spd*. This plasmid was fully assembled only in H1_20d. Contigs co-harboring rep_cluster_1118 and *ant*(*9*)*-*like|pan_4385 were also present in H2_17d and H3_30d. Nevertheless, these contigs were linear, often larger than the fully assembled plasmid and it was not possible to reliably assemble a complete plasmid.

The assembly of complete plasmids from plasmidome samples is a challenging task. We were not able to obtain all completely assembled plasmids. We observed excessive repetitions within contigs, repeated contigs with the same genetic context, and short seemingly circular fragments (<1 kb) which were not truly circular. These issues may be associated with the presence of multiple copies of MGEs, which could lead to a false positive detection of circular fragments in case contigs contain regions flanked by MGEs. The presence of different variants of similar plasmids within a sample could contribute to the occurrence of several repeated contigs with the same genetic content. The assembly failure can also be caused by the chimeric nature of the obtained reads, as observed in the network-based analysis (Fig. 5, Fig. 6) where multiple MOB groups were co-carried on a single read. This could be caused by methodological limitations. The phi29 amplification approach generates tandem repeats of the target plasmid, as confirmed by restriction digestion (41). However, the use of phi29 DNA polymerase is crucial for obtaining sufficient amounts of pDNA.

As we encountered several different issues with assembled genomes, these possibly emerged as an artifact during the assembly process, consistent with recent observations from assembly of metagenomic long-read data from PacBio (42). However, the complex analysis of long-read sequenced plasmidome revealed plasmid-ARGs associations which cannot be analyzed in a short-read metagenomic approach as the sequence depth of the mobilome is not sufficient. The same observations were reported in the previous study where no plasmid-borne ARGs and virulence factors were recovered from metagenome-assembled genomes (32).

### Association of observed results with bacterial species

Among all analyzed samples, the H3_30d was the most distinct from the overall dataset, in terms of its resistome, plasmidome and microbial community.

Taxonomically, H3_30d was characterized by a predominance of Actinobacteria, comprising 69.4% of the total community, whereas Firmicutes, which dominated in all other samples with relative abundances ranging from 80.2% to 99.3%, accounted for only 30.6% in this case (Fig. 7).

**Fig. 7:**
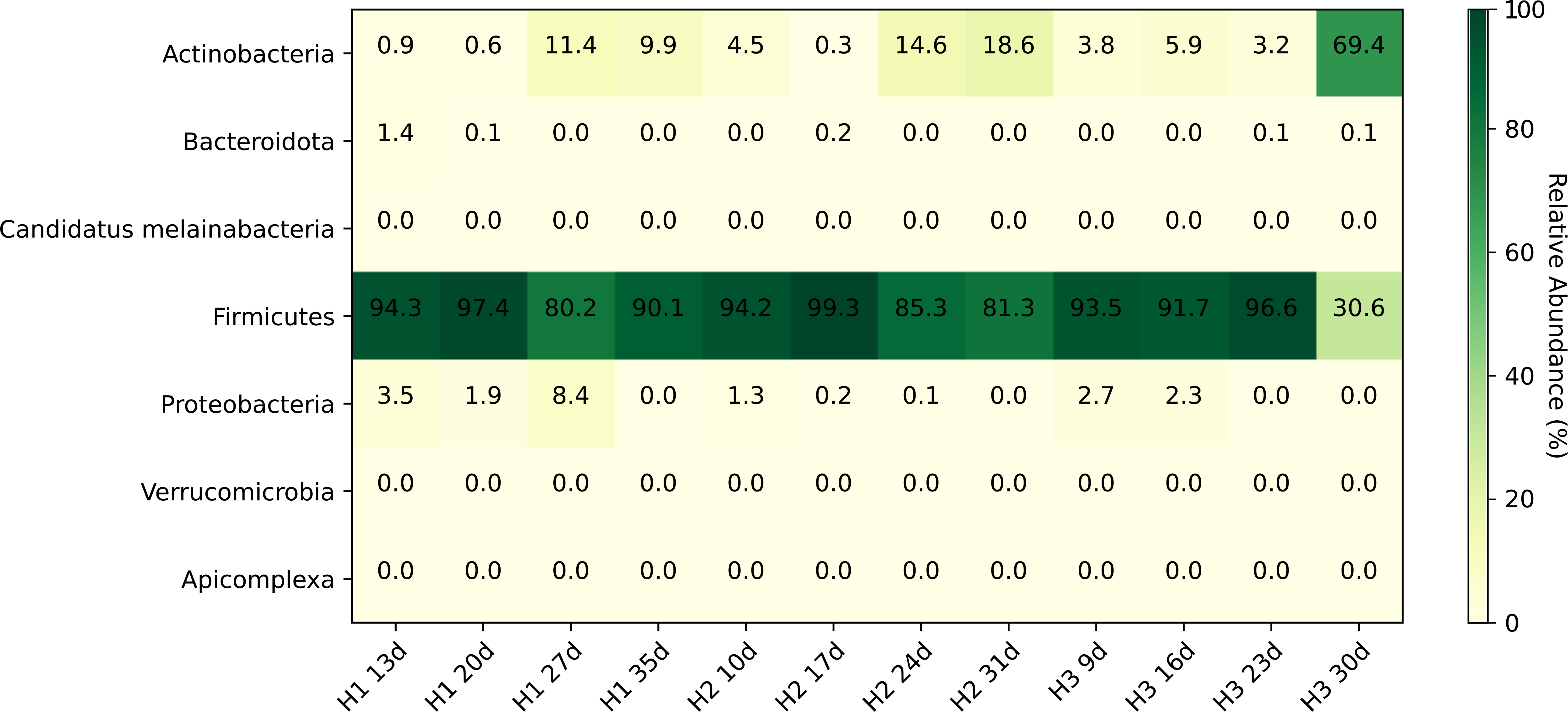
Microbial composition of the samples collected in the chicken farm. Heatmap shows the relative abundance of taxonomic groups (in %) across individual samples based on short-read metagenomic sequencing data.

The presence of Actinobacteria in the H3_30d is likely due to contamination from litter-associated bacteria, as previously reported (43). Given that chickens densely occupy the entire housing area, their feces may contain bacterial remnants from the surrounding litter. This suggests that microbial profiling of fecal samples in such environments should account for potential external bacterial contributions, which may influence downstream analyses and interpretations of gut microbiota composition.

### Technical limitations in plasmidome analysis

This study provides valuable insights into plasmid biology and HGT that would not have been attainable through metagenomic sequencing alone. Unlike standard metagenomic approaches, which are dominated by chromosomal sequences and often fail to detect low-abundance MGEs (32), plasmidome sequencing enabled a more detailed characterization of horizontally transferred ARGs, their associations with specific plasmid types, and their temporal dynamics. Despite the strengths of this study, several methodological limitations should be acknowledged. First, the treatment of extracted pDNA with Plasmid-Safe ATP-dependent DNase, intended to remove chromosomal and linear environmental DNA, may have inadvertently degraded linearized or sheared plasmids. Given that large plasmids are more prone to fragmentation during mechanical lysis and extraction than small plasmids, this step could have contributed to a selective loss of large, low-copy-number plasmids.

The use of phi29 DNA polymerase for plasmid DNA amplification, while necessary to obtain sufficient material for long-read sequencing, may also have introduced a bias towards small, circular plasmids with high copy numbers. This polymerase exhibits preferential amplification of small DNA templates, potentially leading to the overrepresentation of certain plasmid types while obscuring the diversity and abundance of larger or structurally complex plasmids. These findings illustrate the complexity of accurately capturing the plasmidome within metagenomic contexts and point to the importance of continued methodological refinement in plasmidome research.

The processing of sequencing data largely depending on databases of known genes can also represent a limitation of the current research, as well as the use of fixed thresholds (identity ≥ 90%, coverage ≥ 80%) to define gene presence. As a result, novel or highly divergent resistance determinants may remain undetected. However, these limitations are common to many studies relying on sequence-based detection of genes and were recognized in the previous research (44, 45). Despite these challenges, our study benefits from the comprehensive, up-to-date resources; the identification of ARGs was performed using PanRes which integrates multiple major ARGs databases, the plasmid detection was conducted an extensive MOB-suite database, and the reconstructed plasmids were compared to PLSDB, the largest curated repository of complete plasmid sequences.

## Conclusion

Insights into resistance evolution and dissemination require approaches capable of identifying both chromosomal and mobile genetic elements. Although the metagenomic approach provides valuable insights into complex microbiomes, including unculturable bacterial fractions, it provides only limited insight into plasmid interactions. Plasmidome analysis enabled the detection of 133 unique plasmid types across the dataset, compared to only 59 identified by metagenomic sequencing. Plasmidome analysis also suggested that fluoroquinolone resistance genes, that were of particular interest due to the enrofloxacin treatment, were disseminated via MOBP family plasmids, and these were not captured by metagenomic analysis alone. This highlights the added value of plasmid-targeted approaches in resistome surveillance. The associations between plasmids and resistance genes point to dissemination of AMR in the chicken gut microbiome, supporting its role as both a reservoir and an environment favorable for plasmid-mediated gene transfer.

Even despite its technical limitations, targeted plasmidome sequencing offers a complementary perspective, enabling a clearer understanding of gene mobility dynamics and its potential contribution to the persistence and spread of AMR. Integrating both approaches can improve our ability to study resistance in complex microbial ecosystems and to assess the risks associated with HGT in agricultural environments.

## Acknowledgements

We would like to thank Rene Pariza for DNA extraction. We also thank the Center for molecular biology and genetics for providing access to the P2Solo sequencing platform, and Matej Bezdicek for handling the sequencing runs and valuable advice.

## Accession number

Raw sequencing reads from ONT and Illumina sequencing have been deposited in the SRA NCBI under bioproject PRJNA1227156.

## Transparency declarations

None to declare.

## References

1. Simjee S, Ippolito G. 2022. European regulations on prevention use of antimicrobials from january 2022. Rev Bras Med Vet 44.

2. Medicines Agency E. 2023. Sales of veterinary antimicrobial agents in 31 European countries in 2022 - Trends from 2010 to 2022 - Thirteenth ESVAC report 10.2809/766171.

3. Státní veterinární správa ČR. 2019. Národní akční plán pro welfare hospodářských zvířat na období 2020–2025. https://www.svscr.cz/wp-content/files/zvirata/AP_NAP_2019_-_text.pdf. Retrieved 6 May 2025.

4. Roth N, Käsbohrer A, Mayrhofer S, Zitz U, Hofacre C, Domig KJ. 2018. The application of antibiotics in broiler production and the resulting antibiotic resistance in Escherichia coli: A global overview. Poult Sci 98:1791.

5. Unicomb L, Ferguson J, Riley T V., Collignon P. 2003. Fluoroquinolone Resistance in Campylobacter Absent from Isolates, Australia. Emerg Infect Dis 9:1482–1483.

6. Khan I, Bai Y, Zha L, Ullah N, Ullah H, Shah SRH, Sun H, Zhang C. 2021. Mechanism of the Gut Microbiota Colonization Resistance and Enteric Pathogen Infection. Front Cell Infect Microbiol 11:716299.

7. Klimien I, Virgailis M, Kerzien S, Siug R, Mockeli Unas R, Modestas |, Zauskas R. Evaluation of genotypical antimicrobial resistance in ESBL producing Escherichia coli phylogenetic groups isolated from retail poultry meat 10.1111/jfs.12370.

8. Błażejewska A, Zalewska M, Grudniak A, Popowska M. 2022. A Comprehensive Study of the Microbiome, Resistome, and Physical and Chemical Characteristics of Chicken Waste from Intensive Farms. Biomolecules 12:1132.

9. Harrison E, Brockhurst MA. 2012. Plasmid-mediated horizontal gene transfer is a coevolutionary process. Trends Microbiol 20:262–267.

10. Lipworth S, Matlock W, Shaw L, Vihta KD, Rodger G, Chau K, Barker L, George S, Kavanagh J, Davies T, Vaughan A, Andersson M, Jeffery K, Oakley S, Morgan M, Hopkins S, Peto T, Crook D, Walker AS, Stoesser N. 2024. The plasmidome associated with Gram-negative bloodstream infections: A large-scale observational study using complete plasmid assemblies. Nature Communications 2024 15:1 15:1–11.

11. Brown Kav A, Benhar I, Mizrahi I. 2013. A method for purifying high quality and high yield plasmid DNA for metagenomic and deep sequencing approaches. J Microbiol Methods 95:272–279.

12. Kirstahler P, Teudt F, Otani S, Aarestrup FM, Pamp SJ. 2021. A Peek into the Plasmidome of Global Sewage. mSystems 6.

13. Schlüter A, Krause L, Szczepanowski R, Goesmann A, Pühler A. 2008. Genetic diversity and composition of a plasmid metagenome from a wastewater treatment plant. J Biotechnol 136:65–76.

14. Li AD, Li LG, Zhang T. 2015. Exploring antibiotic resistance genes and metal resistance genes in plasmid metagenomes from wastewater treatment plants. Front Microbiol 6:154890.

15. Che Y, Xia Y, Liu L, Li AD, Yang Y, Zhang T. 2019. Mobile antibiotic resistome in wastewater treatment plants revealed by Nanopore metagenomic sequencing. Microbiome 7:1–13.

16. Jørgensen T, Sparholt ;, Hansen M, Asser ;, Xu Z;, Tabak MA;, Sørensen S, Johannes ;, Hansen LH. 2017. Plasmids, viruses, and other circular elements in rat gut 10.1101/143420.

17. Kav AB, Sasson G, Jami E, Doron-Faigenboim A, Benhar I, Mizrahi I. 2012. Insights into the bovine rumen plasmidome. Proc Natl Acad Sci U S A 109:5452–5457.

18. Fang Z, Tan J, Wu S, Li M, Xu C, Xie Z, Zhu H. 2019. PPR-Meta: a tool for identifying phages and plasmids from metagenomic fragments using deep learning. Gigascience 8:1–14.

19. Hunter JD. 2007. Matplotlib: A 2D graphics environment. Comput Sci Eng 9:90–95.

20. Clausen PTLC, Aarestrup FM, Lund O. 2018. Rapid and precise alignment of raw reads against redundant databases with KMA. BMC Bioinformatics 19:1–8.

21. Martiny H-M, Pyrounakis N, Lukjančenko O, Petersen TN, Aarestrup FM, Clausen PTLC, Munk P. PanRes - Collection of antimicrobial resistance genes 10.5281/ZENODO.10091602.

22. Johansson MHK, Bortolaia V, Tansirichaiya S, Aarestrup FM, Roberts AP, Petersen TN. 2021. Detection of mobile genetic elements associated with antibiotic resistance in Salmonella enterica using a newly developed web tool: MobileElementFinder. Journal of Antimicrobial Chemotherapy 76:101–109.

23. Robertson J, Nash JHE. 2018. MOB-suite: software tools for clustering, reconstruction and typing of plasmids from draft assemblies. Microb Genom 4:e000206.

24. Schwarzerova J, Labanava A, Rychlik I, Varga M, Cejkova D. 2023. A minireview on the bioinformatics analysis of mobile gene elements in microbiome research. Frontiers in Bacteriology 2:1275910.

25. Shannon P, Markiel A, Ozier O, Baliga NS, Wang JT, Ramage D, Amin N, Schwikowski B, Ideker T. 2003. Cytoscape: A Software Environment for Integrated Models of Biomolecular Interaction Networks. Genome Res 13:2498.

26. Molano LAG, Hirsch P, Hannig M, Müller R, Keller A. 2025. The PLSDB 2025 update: enhanced annotations and improved functionality for comprehensive plasmid research. Nucleic Acids Res 53:D189–D196.

27. Schwengers O, Jelonek L, Dieckmann MA, Beyvers S, Blom J, Goesmann A. 2021. Bakta: rapid and standardized annotation of bacterial genomes via alignment-free sequence identification. Microb Genom 7:685.

28. Blanco-Míguez A, Beghini F, Cumbo F, McIver LJ, Thompson KN, Zolfo M, Manghi P, Dubois L, Huang KD, Thomas AM, Nickols WA, Piccinno G, Piperni E, Punčochář M, Valles-Colomer M, Tett A, Giordano F, Davies R, Wolf J, Berry SE, Spector TD, Franzosa EA, Pasolli E, Asnicar F, Huttenhower C, Segata N. 2023. Extending and improving metagenomic taxonomic profiling with uncharacterized species using MetaPhlAn 4. Nat Biotechnol 41:1633–1644.

29. Parks DH, Chuvochina M, Rinke C, Mussig AJ, Chaumeil PA, Hugenholtz P. 2022. GTDB: an ongoing census of bacterial and archaeal diversity through a phylogenetically consistent, rank normalized and complete genome-based taxonomy. Nucleic Acids Res 50:D785–D794.

30. Josefsen MH, Andersen SC, Christensen J, Hoorfar J. 2015. Microbial food safety: Potential of DNA extraction methods for use in diagnostic metagenomics. J Microbiol Methods 114:30–34.

31. Juricova H, Matiasovicova J, Kubasova T, Cejkova D, Rychlik I. 2021. The distribution of antibiotic resistance genes in chicken gut microbiota commensals. Sci Rep 11:1–10.

32. Maguire F, Jia B, Gray KL, Lau WYV, Beiko RG, Brinkman FSL. 2020. Metagenome-assembled genome binning methods with short reads disproportionately fail for plasmids and genomic islands. Microb Genom 6:1–12.

33. Osorio V, Sabater i Mezquita A, Balcázar JL. 2023. Comparative metagenomics reveals poultry and swine farming are hotspots for multidrug and tetracycline resistance. Environmental Pollution 322:121239.

34. Hegde N V., Kariyawasam S, DebRoy C. 2016. Comparison of antimicrobial resistant genes in chicken gut microbiome grown on organic and conventional diet. Vet Anim Sci 1–2:9–14.

35. Xiong W, Wang Y, Sun Y, Ma L, Zeng Q, Jiang X, Li A, Zeng Z, Zhang T. 2018. Antibiotic-mediated changes in the fecal microbiome of broiler chickens define the incidence of antibiotic resistance genes. Microbiome 6:1–11.

36. Delsol AA, Sunderland J, Woodward MJ, Pumbwe L, Piddock LJV, Roe JM. 2004. Emergence of fluoroquinolone resistance in the native Campylobacter coli population of pigs exposed to enrofloxacin. Journal of Antimicrobial Chemotherapy 53:872–874.

37. Sato T, Ito R, Kawamura M, Fujimura S. 2022. The Risk of Emerging Resistance to Trimethoprim/ Sulfamethoxazole in Staphylococcus aureus. Infect Drug Resist 15:4779–4784.

38. Mazurek J, Bok E, Stosik M, Baldy-Chudzik K. 2015. Antimicrobial Resistance in Commensal Escherichia coli from Pigs during Metaphylactic Trimethoprim and Sulfamethoxazole Treatment and in the Post-Exposure Period. Int J Environ Res Public Health 12:2150.

39. Wang Y, Dagan T. 2024. The evolution of antibiotic resistance islands occurs within the framework of plasmid lineages. Nat Commun 15:1–13.

40. Duxbury SJN, Arjan J, De Visser GM, Alderliesten JB, Zwart MP, Stegeman A, Fischer EAJ. 2021. Chicken gut microbiome members limit the spread of an antimicrobial resistance plasmid in Escherichia coli 10.1098/rspb.2021.2027.

41. Dean FB, Nelson JR, Giesler TL, Lasken RS. 2001. Rapid Amplification of Plasmid and Phage DNA Using Phi29 DNA Polymerase and Multiply-Primed Rolling Circle Amplification. Genome Res 11:1095–1099.

42. Trigodet F, Sachdeva R, Banfield JF, Eren AM. 2025. Assemblies of long-read metagenomes suffer from diverse errors. bioRxiv 2025.04.22.649783.

43. Rychlik I, Karasova D, Crhanova M. 2023. Microbiota of Chickens and Their Environment in Commercial Production. Avian Dis 67:1–9.

44. Papp M, Solymosi N. 2022. Review and Comparison of Antimicrobial Resistance Gene Databases. Antibiotics 11:339.

45. Gschwind R, Ugarcina Perovic S, Weiss M, Petitjean M, Lao J, Coelho LP, Ruppé E. 2023. ResFinderFG v2.0: a database of antibiotic resistance genes obtained by functional metagenomics. Nucleic Acids Res 51:W493.

